# Electrical signature of heterogeneous human mesenchymal stem cells

**DOI:** 10.1101/2021.11.08.467665

**Authors:** Tunglin Tsai, Prema D. Vyas, Lexi L. Crowell, Mary Tran, Destiney W. Ward, Yufan Qin, Angie Castro, Tayloria N.G. Adams

## Abstract

Human mesenchymal stem cells (hMSCs) have gained traction in transplantation therapy due to their immunomodulatory, paracrine, immune-evasive, and multipotent differentiation potential. Given the heterogeneous nature of hMSCs, therapeutic treatments and robust in vivo and in vitro experiments require additional biomarkers to ensure reproducibility when using these stem cells. In this work, we utilized dielectrophoresis (DEP), a label-free electrokinetic phenomenon, to investigate and quantify the heterogeneity of hMSCs derived from the bone marrow (BM) and adipose tissue (AD). Through computer simulation, we identified that the transient slope of the DEP force spectra can be used as a metric of heterogeneity. The electrical properties of BM-hMSCs were compared to homogeneous mouse fibroblasts (NIH-3T3), human fibroblasts (WS1), and human embryonic kidney cells (HEK-293). BM-hMSCs DEP profile was most different from HEK-293 cells. We compared the DEP profiles of BM-hMSCs and AD-hMSCs and found they have similar membrane capacitances, differing cytoplasm conductivity, and transient slopes. Inducing both populations to differentiate into adipocyte and osteocyte cells revealed they behave differently in response to differentiation-inducing cytokines. Histology and RT-qPCR analyses of the differentiation-related genes revealed differences in heterogeneity between BM-hMSCs and AD-hMSCs. The differentiation profiles correlate well with the DEP profiles developed and indicate that these BM-hMSCs have higher differentiation potential than AD-hMSCs. Our results demonstrate using DEP, membrane capacitance, cytoplasm conductivity, and transient slope can uniquely characterize the inherent heterogeneity of hMSCs to guide robust and reproducible stem cell transplantation therapies.

## INTRODUCTION

Despite recent advances in stem cell therapies, the inherent heterogeneity of adult stem cell populations remains a challenge to improving their therapeutic efficacy. Human mesenchymal stem cells (hMSCs) possess regenerative properties because they secrete bioactive molecules beneficial for cell survival and function, they support the immune system, and differentiate into multiple cell types^[1, 2]^. HMSCs heterogeneity has been attributed to the varied outcomes observed in clinical trials ^[3]^. For instance, studies treating spinal cord injury demonstrated that patients who received bone marrow-derived (BM) hMSCs had improvements in muscle spasms ^[4]^, and those who received umbilical cord-derived (UC) hMSCs had restored spinal cord function ^[5]^. In these stem cell therapies, the heterogeneity of BM-hMSCs and UC-hMSCs impacted patient outcomes. One method proven successful for reducing the heterogeneity of stem cells is the implementation of dielectrophoresis (DEP) on microfluidic devices ^[6]^. With this technique, important gains were made in characterizing and generating purified subsets of astrocyte-biased^[6]^ and neuron-biased^[7]^ cells from neural stem and progenitor cells (NSPCs). Thus, DEP holds promise as a cell analysis tool for characterizing, understanding, and eventually reducing the heterogeneity of hMSCs.

The heterogeneity of hMSCs can be classified by their morphology, cell size, electrical properties, cell fate, gene expression, and surface protein expression. Haasters et al.^[8]^, identified three morphologically and functionally distinct subtypes of hMSCs, rapidly self-renewing (RS) cells, spindle-shaped (SS) cells, and slowly-replicating flattened cells (FC). Larger cells, the FCs, had slow motility and more senescent cells while smaller cells, the RSs, had fast motility and less senescent cells. Cell size and motility were ranked as RS < SS < FC and larger cells had osteocyte attributes ^[8]^. Yang et al.^[9]^ found corresponding morphology results but also reported a dependence of morphology on passage number (i.e., in vitro age). Early passage hMSCs maintained SS morphology while later passages exhibited irregular FC morphology and larger size. These data emphasize that cell size, morphology, and in vitro age are linked to heterogeneity.

HMSCs generate mature adipocytes, chondrocytes, osteocytes, myocytes, fibroblasts, and hepatocytes^[1, 2]^, which contributes their inherent heterogeneity. Impedance measurements have been reported to detect hMSC osteogenic and adipogenic differentiation ^[10, 11]^, serving as an indicator of heterogeneity. HMSCs cell fate, gene, and surface protein expression vary based on sources^[12–14]^, cell culture methods^[9, 15]^, and in vitro age^[9]^. Mohamed-Ahmed et al.^[13]^, examined BM-hMSCs and adipose tissue (AD) derived hMSCs from multiple donors. Even with donor matching (i.e., BM-hMSC and AD-hMSCs from the same person), there were differences in osteogenic, adipogenic, and chondrogenic cell fate potentials. The BM-hMSCs had higher osteogenic and chondrogenic potential and AD-hMSCs had higher adipogenic potential. Lee et al.^[16]^ found that BM-hMSCs and dental pulp-derived hMSCs had similar cell morphology, surface protein expression, and differentiation potential. Many hMSCs heterogeneity assessments require differentiation and antibodies, label-based biomarkers. In order to select the proper antibodies prior knowledge of the cell sample’s heterogeneity is required. In many cases the antibodies used overlap with other cell types^[17, 18]^. Another disadvantage is that the incorporation of antibodies increases cell processing steps and these techniques are expensive^[19]^. Therefore, having reliable biomarkers that remove the requirement of antibodies for assessing and understanding hMSCs heterogeneity is desirable.

Membrane capacitance, a label-free biomarker, has arisen as a good indicator of stem cell heterogeneity^[20, 21]^, and is a measure of a cell’s ability to store charge. Since hMSCs are heterogeneous containing different cells, each cell type will have a unique membrane capacitance value. DEP is a cell analysis technique that uses nonuniform electric fields and the presence of ions to measure the membrane capacitance of cells. DEP provides bulk measurements of a cell population’s membrane capacitance using cell polarization at select frequencies. Adams et al.^[20]^ used DEP to characterize the membrane capacitance and membrane permittivity of BM-hMSCs. They found there were at least two different cell types present over a wide range of frequencies. Similarly, Giduthur et al.^[22]^, found that murine MSCs were heterogeneous in their DEP response and cell morphology with at least three different cells types present. Additionally, DEP has been used to examine the heterogeneity of human NSPCs^[6, 21]^, mouse NSPCs^[23]^ and red blood cells^[24]^.

Correlating biological assessments of heterogeneity to engineering assessments is important. In this study, we tested whether the heterogeneity of BM-hMSCs and AD-hMSCs with potential clinical relevance could be quantified using DEP. We utilized two different DEP-based microfluidic devices, quadrapole and 3DEP, to examine hMSCs polarizability and DEP response with homogeneous mouse fibroblasts (NIH-3T3), human skin fibroblasts (WS1), and human kidney (HEK-293) cell populations. Cell motion was imaged and analyzed using multiple measures: relative DEP force, transient slope, membrane capacitance, and cytoplasm conductivity. Cell size was also quantified as a measure of heterogeneity. hMSCs heterogeneity was verified with adipogenic and osteogenic differentiation, histological staining, and RT-qPCR analysis of differentiation-related genes. We found that BM-hMSCs have distinct electrical signatures compared to NIH-3T3, WS1, and HEK-293 cells. Additionally, our data shows that BM-hMSCs and AD-hMSCs have similar membrane capacitance value but unique cytoplasm conductivity and transient slope values. Differentiation results showed that both BM-hMSCs and AD-hMSCs change into adipocytes and osteocytes to varying degrees.

### Dielectrophoresis (DEP) Theory

DEP employs nonuniform electric fields, created with electrodes, to induce cell movement based on the polarizability and inherent electrical properties (capacitance and conductivity) of cells^[25]^. For DEP polarization to occur, cells are placed in a conductive medium. When the alternating current electric field, defined by a waveform, voltage and frequency, is applied the field interacts with ions available in the medium causing them to move and align around the cell (i.e., polarization)^[26]^, which is directly impacted by the cell’s electrical properties. The induced DEP force that causes cell movement is given by 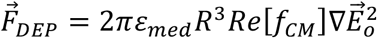 ^[25]^ where *ε_med_* is the medium permittivity (F/m), *R* is the cell radius (μm), *Re[f_CM_*] is the real part of the Clausius-Mossotti factor (unitless), and 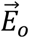 is the electric field (V/m). The *f_CM_* describes cell motion in the electric field and is defined as^[25]^,

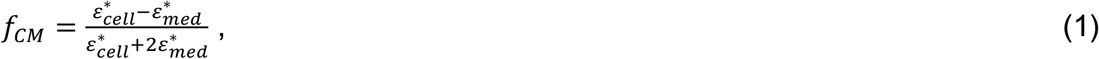

where, 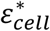 is the complex permittivity of the cell and 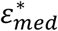 is the complex permittivity of the suspending medium. Cells are modeled as core-shell particles such that 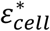 gets replaced with an effective complex permittivity, 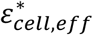, defined as^[25]^,

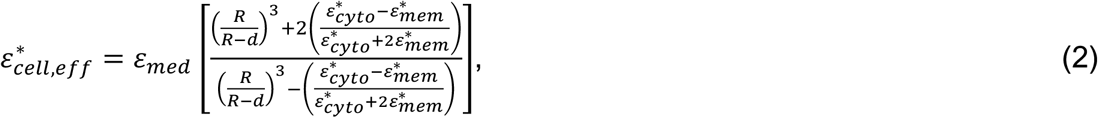

where, *d* is the plasma membrane thickness (μm), 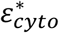 is the complex permittivity of the cell cytoplasm (unitless), and 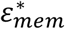 is the complex permittivity of the cell membrane (unitless). The complex permittivity is given by^[25]^,

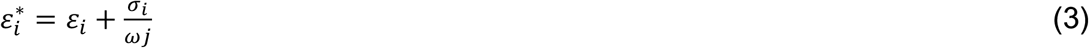

where *ε_i_* is the permittivity, *σ_i_* is the conductivity, and *i* = cell, cytoplasm, membrane, or medium. Cell motion in response to the applied electric field can be used to determine sample heterogeneity.

Cells have distinct frequency-dependent polarizability that can be used for identification with DEP. Polarized cells will exhibit either positive DEP force, wherein cells move to areas of high electric field strength, or negative DEP force, wherein cells move to areas of low electric field strength^[25]^. At radio frequencies, 100 kHz to 10 MHz, cell polarizability is affected by their plasma membrane and cytoplasm. At lower radio frequencies, cells are polarized, and the resistance of the cell membrane shields the interior components of the cell (i.e., cytoplasm, mitochondria, nucleus, etc.) from the external electric field probing structural (blebs, folding, etc.) and molecular features (proteins) of the cell membrane ^[27]^. At higher radio frequencies, cells are polarized, and the electric field penetrates the cell surface, analyzing the interior cell components^[27]^. Thus, low and high frequency DEP measurements provide information about the cell membrane and cytoplasm, respectively.

Cell DEP polarization is characterized experimentally by measuring the positive and negative DEP responses at specific frequencies. Visually, this means that at specific frequencies cells will appear along electrode edges for positive DEP and not along electrode edges for negative DEP, which is used to build a DEP response spectrum. With heterogeneous cell populations, both negative and positive DEP cell behaviors are visualized at a single frequency. Cells have two characteristic crossover frequencies (*f_xo_*), meaning there is no net movement in response to the electric field, influenced by cell structural and interior components. The low *f_xo_* is determined by cell size, shape, and plasma membrane and the higher *f_xo_* is influenced by the cytoplasm. Using data points from the DEP response spectra the electrical properties of the cell membrane (capacitance and conductivity) and cytoplasm (conductivity and permittivity) are estimated. Membrane capacitance, *C_mem_* (mF/m^2^), is a function of *f_xo_* and given as 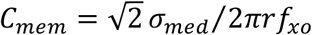 ^[25]^ where *σ_med_* is the medium conductivity (μS/cm) and r is the cell radius (μm). Membrane permittivity, *ε_mem_* (unitless), is proportional to *C_mem_* and given as ^[20]^

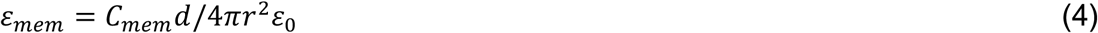

where ε_0_ = 8.85 × 10 ^−12^ is the vacuum permittivity. The cytoplasm conductivity and permittivity are estimated by Eq. 3.

## MATERIALS AND METHODS

### Cell culture

BM-hMSCs (ATCC, Manassas, VA) were cultured in MSC basal media supplemented with 7% fetal bovine serum (FBS) (ATCC, Manassas, VA), 15 ng/mL IGF-1 (ATCC, Manassas, VA), 125 pg/mL FGF-2 (ATCC, Manassas, VA), 2.4mM L-Alanyl-L-Glutamine (ATCC, Manassas, VA), and 0.1X antibiotic-antimycotic (Life Technologies, Carlsbad, CA). AD-hMSC (ATCC, Manassas, VA) were cultured in MSC basal media supplemented with 2% FBS, 5 ng/mL FGF-1 (ATCC, Manassas, VA), 5 ng/mL FGF-2, 5 ng/mL EGF (ATCC, Manassas, VA), and 0.1X antibiotic-antimycotic. Both BM-hMSC and AD-hMSC were grown in tissue culture-treated T-75 flasks by seeding 5,000 cells/cm^2^ and passaged at ~80% confluence.

During cell passaging, the growth medium was aspirated, and the monolayer was rinsed once using 1X DPBS (Life Technologies, Carlsbad, CA). The cells were dissociated using 0.05% Trypsin-EDTA (Life Technologies, Carlsbad, CA) for 4 min at 37°C. When most of the cells were dissociated from the growth surface, residual trypsin activity was neutralized with an equal volume of a neutralizing solution consisting of 5% FBS in 1X DPBS. The removed cells were centrifuged at 275 x g and resuspended in the new growth medium and counted for accurate seeding.

NIH-3T3 cells (ATCC, Manassas, VA) were maintained in Dulbecco’s Modified Eagle Medium supplemented with 10% newborn calf serum. WS1 and HEK-293 cells were obtained from ATCC (Manassas, VA) and cultured in Eagle’s Minimum Essential Medium (Life Technologies, Carlsbad, CA) supplemented with 10% FBS. All cells were seeded at 5,000 cells/cm^2^ and passaged at ~80% confluence.

### DEP characterization

On the day of the experiment, cells were trypsinized and resuspended in an isotonic DEP buffer solution consisting of 8.5% (w/v) sucrose and 0.3% (w/v) glucose.

The conductivity was adjusted to 100 μS/cm with RPMI-1640 medium. The cells were washed 3 times before resuspending in DEP buffer at 1 × 10^6^ cells/mL. First, a visual DEP screening of hMSC heterogeneity was completed with a quadrupole device (fabricated using previously published techniques^[6, 23]^; electrode dimensions: 50 μm wide with 200 μm spacing). At select frequencies and 10 Vpp cell movement was recorded.

A full DEP characterization was performed using the 3DEP analyzer (LabTech, East Sussex, UK). The 3DEP chip was primed with 70% ethanol, rinsed 3 times with 100 μL Milli-Q water, and another 3 times with 100 μL of DEP buffer. Using a beveled 200 μL pipet tip, 80 μL of the resuspended cells were introduced into the chip and the chip was covered with an 18 x 18 mm glass coverslip. The loaded chip was run on the 3DEP analyzer using 20 log-linear frequencies ranging from 2 kHz to 20 MHz at 10 Vpp for 60 s. The DEP response was evaluated using the 3DEP analyzer software and individual runs with an R-value > 0.9 were kept for further analysis (~10 runs for each independent experiment). For all experiments, the cells were imaged on a hemacytometer, and cell–size measured using ImageJ software.

### MATLAB modeling of hMSCs electrical properties

The 3DEP analyzer outputs the relative DEP response based on changes in light intensity as a function of frequency^[28]^. The intensity data for each independent DEP experiment were pooled together for a quartile test and identified outliers were removed at specific frequencies. Next, these data (light intensity vs frequency) were imported into MATLAB. The core-shell spherical DEP polarization model was fitted to the experimental data using the fminsearchbnd function, which implements non-linear fitting^[29]^. The *σ_mem_*, *ε_mem_*, *σ_cyto_*, *ε_cyto_*, and a linear scalar were simultaneously and iteratively adjusted to give the highest R-value. The linear scalar was used to transform the DEP response into the relative DEP force spectrum as a function of frequency. Cell size and conductivity of the DEP buffer were fixed model parameters. The *C_mem_* was estimated using Eq. (4) assuming 10 nm for cell membrane thickness. This modeling of hMSCs’ electrical properties was carried out two ways: non-linear fitting to each individual DEP runs (fitting to 20 data points) and non-linear fitting to pooled data (fitting to 200 data points). Finally, the transient slope of the relative DEP force spectrum was calculated by fitting a linear trendline to 20%-80% of the rise time (adapted from^[30]^).

### HMSCs differentiation

Tissue culture-treated 6-well plates were coated with 0.2% gelatin to prepare for hMSC differentiation. For coating, lyophilized porcine skin gelatin (Sigma Aldrich, St. Louis, MO) was dissolved in Milli-Q water and sterilized by autoclaving. 900 μL of the gelatin solution was added to the 6-well plate and allowed to coat at room temperature for 30 min. The excess gelatin solution was aspirated, and the plates were placed in the biosafety cabinet to dry for at least 2 h.

Both the BM-hMSCs and AD-hMSCs were seeded at 10,000 cells/cm^2^ in the 6-well plate and proliferated for 2 days before inducing differentiation. For osteocyte differentiation, the growth medium was aspirated and rinsed once with 1X DPBS and replaced with an osteogenic medium, which consisted of αMEM (Life Technologies, Carlsbad, CA) supplemented with 10% FBS, 50 μg/ml l-ascorbic acid 2-phosphate (FUJIFILM Wako, Richmond, VA), 100 nM dexamethasone (MP Biomedicals, Irvine, CA), 10 mM β-glycerophosphate (Alfa Aesar, Haverhill, MA), and 0.1X antibiotic-antimycotic. For adipocyte differentiation, the growth medium was replaced with StemPro adipogenesis media (Life Technologies, Carlsbad, CA). The differentiation media was changed every 4 days during the 21-day differentiation. Care was taken to make sure the monolayer of differentiating cells was not disturbed or exposed to air.

### Histology

Differentiated hMSCs were fixed with 4% paraformaldehyde for 15 min at room temperature. After fixing, hMSCs were rinsed 3 times with 1X DPBS and stored in 0.05% sodium azide in 1x PBS at 4°C. Alizarin Red S (ARS) staining of calcium deposits was used to assess osteocyte differentiation. An ARS stock solution was made by dissolving 40mM ARS in Milli-Q water and sterile filtered. HMSCs were incubated in ARS for 15 min at room temperature. The ARS was aspirated, and the cells were rinsed with Milli-Q water until all unbounded ARS was removed. Oil Red O (ORO) staining of lipid accumulation was used to assess adipocyte differentiation. The ORO stock solution was prepared by mixing 0.5% (w/v) ORO dissolved in pure isopropanol with Milli-Q water at a 3:2 ratio. The solution sat for 20 min and was sterile filtered. HMSCs were incubated in ORO for 10 min at room temperature. ORO was aspirated and the cells were rinsed 3 times with Milli-Q water. All cell samples were imaged immediately after staining using EVOS XL (ThermoFisher, Frederick, MD).

### Gene expression

Reverse transcription-quantitative polymerase chain reaction (RT-qPCR) was used to quantify the gene expression changes during hMSC differentiation. RNA was extracted using an RNA micro prep kit (Zymo Research, Irvine, CA). The differentiation media was aspirated, and RNA was liberated from the monolayer by adding the lysis buffer and purified following the manufacturer’s protocol. The cDNA was synthesized using Lunascript reverse transcription master mix (New England Biolabs, Ipswich, MA). Collagen type 1 alpha 1 (COL1A1), alkaline phosphatase (ALPL), and runt-related transcription factor 2 (RUNX2) were used to quantify osteogenesis; and adiponectin (ADIPOQ), fatty acid binding protein 4 (FABP4), and peroxisome proliferator activated receptor gamma (PPARG) were used to quantify adipogenesis. Relative quantification of mRNA expression was obtained using ΔΔCt method using glyceraldehyde-3-phosphate dehydrogenase (GAPDH) as the endogenous control. Primer sequences are in supplemental Table S1.

### Statistical analysis

Statistical analysis for DEP measurements was completed using One-Way ANOVA with Tukey post hoc test for multiple comparisons. For gene expression profiles statistical analysis was completed using two-tailed unpaired student T test with Fisher’s LSD post hoc test. Biological replicates are listed as “n” in figure legends.

## RESULTS AND DISCUSSION

### DEP heterogeneity assessment of hMSCs compared to homogeneous cell populations

One benefit to DEP cell analysis is the generation of images of cells responding to the electric field. An indicator of heterogeneity is the visualization of cells displaying positive and negative DEP at a single frequency. We previously found that the heterogeneity of BM-hMSCs is visible at frequencies above 1 MHz^[20]^. Thus, we screened the DEP response of BM-hMSCs from 2 kHz to 20 MHz, to capture the lower *f_xo_* and approach the higher *f_xo_*. In Figure 1A, with the electric field applied at 9 kHz, the heterogeneity of BM-hMSCs is observed in the top image and annotated in the bottom image for cells experiencing positive DEP (blue circles) and negative DEP (green circles). Next, a MATLAB function was written to simulate the heterogeneity of hypothetical cell populations. First, the relative DEP force curve was generated for three homogeneous cells populations, dashed lines in Figure 1B, with *C_mem_* values of 10, 30, and 75 mF/m^2^ while holding other parameters constant (*ε_cyto_,σ_cyto_,σ_mem_,σ_med_*,d)o To mimic hypothetical heterogeneous cell populations numerous DEP curves were generated and averaged by ranging *C_mem_* from 15 to 45 mF/m^2^ and 10 to 75 mF/m^2^ for low and high heterogeneity, respectively (Figure 1B solid lines). All curves were modeled using Eqs. (1)–(3). The transient slope, Figure 1C, was determined by fitting a linear trendline to the DEP force curve. Gradual slopes are characteristic of heterogeneity and steeper slopes are characteristic of homogeneity. We will use transient slope as a metric to quantify the heterogeneity of hMSCs.

**Figure 1.**
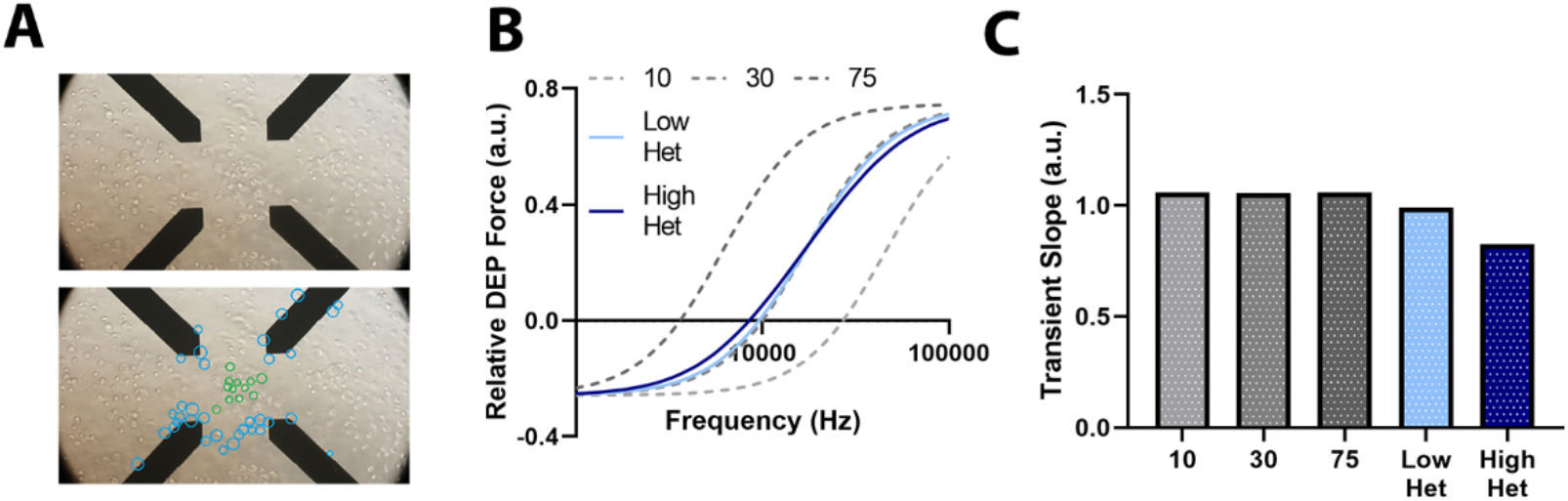
Illustration of cellular heterogeneity using DEP. (A) Images of BM-hMSCs’ DEP responses at 9 kHz for 30 s in the electric field. Bottom image annotates cells experiencing positive and negative DEP (blue, green circles). (B) MATLAB simulation of the DEP force of hypothetical homogeneous (*C_mem_* = 10, 30, and 75 mF/m^2^), low heterogeneous (*C_mem_* = 15-45 mF/m^2^), and high heterogeneous cell populations (*C_mem_* = 10-75 mF/m^2^). (C) Transient slope measured from the DEP force curves in (B).

HMSCs’ heterogeneity was assessed using DEP and Figure 2 outlines the experimental workflow. First, the electrical properties of BM-hMSCs were compared to control homogeneous cell populations: HEK-293, NIH-3T3, and WS1. Then, BM-hMSCs and AD-hMSCs were compared. Approximately 2 × 10^6^ cells were prepared for characterization in DEP buffer solution, Figure 2A. The cells were tested with the quadrapole and 3DEP devices, Figure 2B. The function of BM- and AD-hMSCs was assessed by inducing adipocyte and osteocyte differentiation, Figure 2C. The DEP force spectra of cells from multiple technical replicate measurements were combined and the electrical properties of all cells were extracted using equations in Eqs. (1)–(3), Figure 2D.

**Figure 2.**
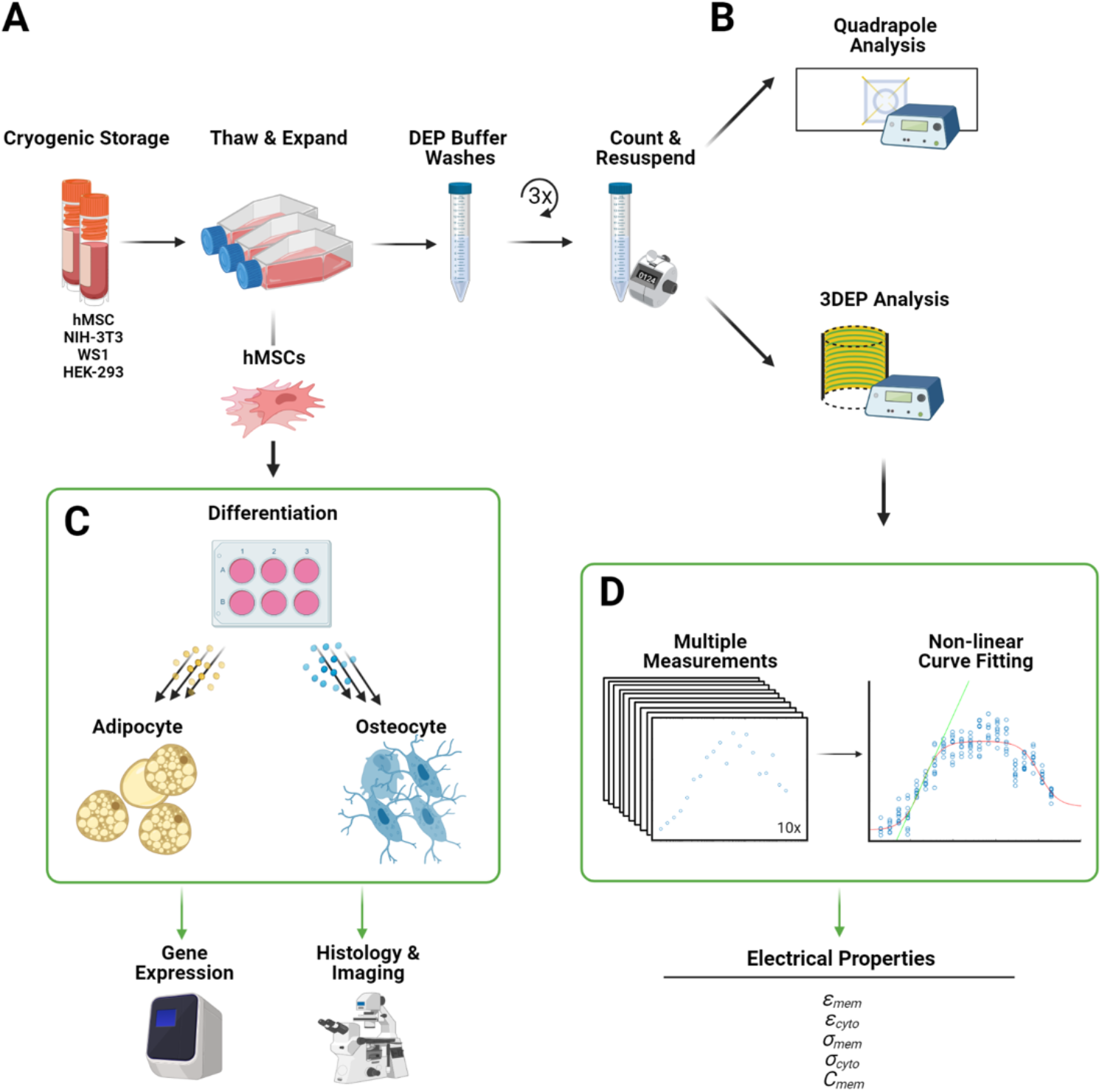
DEP experimental workflow for characterizing heterogeneity. (A) hMSCs, HEK-293, NIH-3T3, and WS1 cells were obtained from cryogenic storage, thawed, and expanded in proliferation media. About 2 × 10^6^ cells were prepared in DEP buffer solution for (B) Quadrapole or 3DEP analysis. For the hMSCs an additional 3 × 10^6^ cells were prepared for (C) adipocyte and osteocyte differentiation. (D) The DEP force spectra obtained from (B) were used to estimate the electrical properties of all cell types. Figure created with BioRender.com.

The DEP force spectra of BM-hMSCs and homogenous controls are shown in Figure 3A. At frequencies below 10 kHz, all cells experience negative DEP force, and the HEK-293 cells experience the strongest negative DEP. At frequencies 10-100 kHz all the cells experience positive DEP, and HEK-293 remain distinguished from the other cells. From 100 kHz to 20 MHz, all cells experience positive DEP and are not discernible. The error bars represent the standard deviation; some are too small for the bars to be visible. Average cell radius is given in Figure 3B, the BM-hMSCs have the largest radius 8.59 *μm* and WS1 having the smallest radius 7.28 μm; the size of each cell is statistically significant. In Figure 3C–3H, we performed non-linear fitting to each individual DEP measurement in order to estimate the cell electrical properties (solid green circles). The MATLAB function, same used in Figure 1, yielded sub-optimal results for *ε_cyto_* (reached upper limit, Figure 3E) and *σ_mem_* (reached its lower limit, Figure 3H). This was resolved by pooling the DEP measurements for 10 runs then performing the non-linear data fitting, open black circles are technical replicates and bars are the total average.

**Figure 3.**
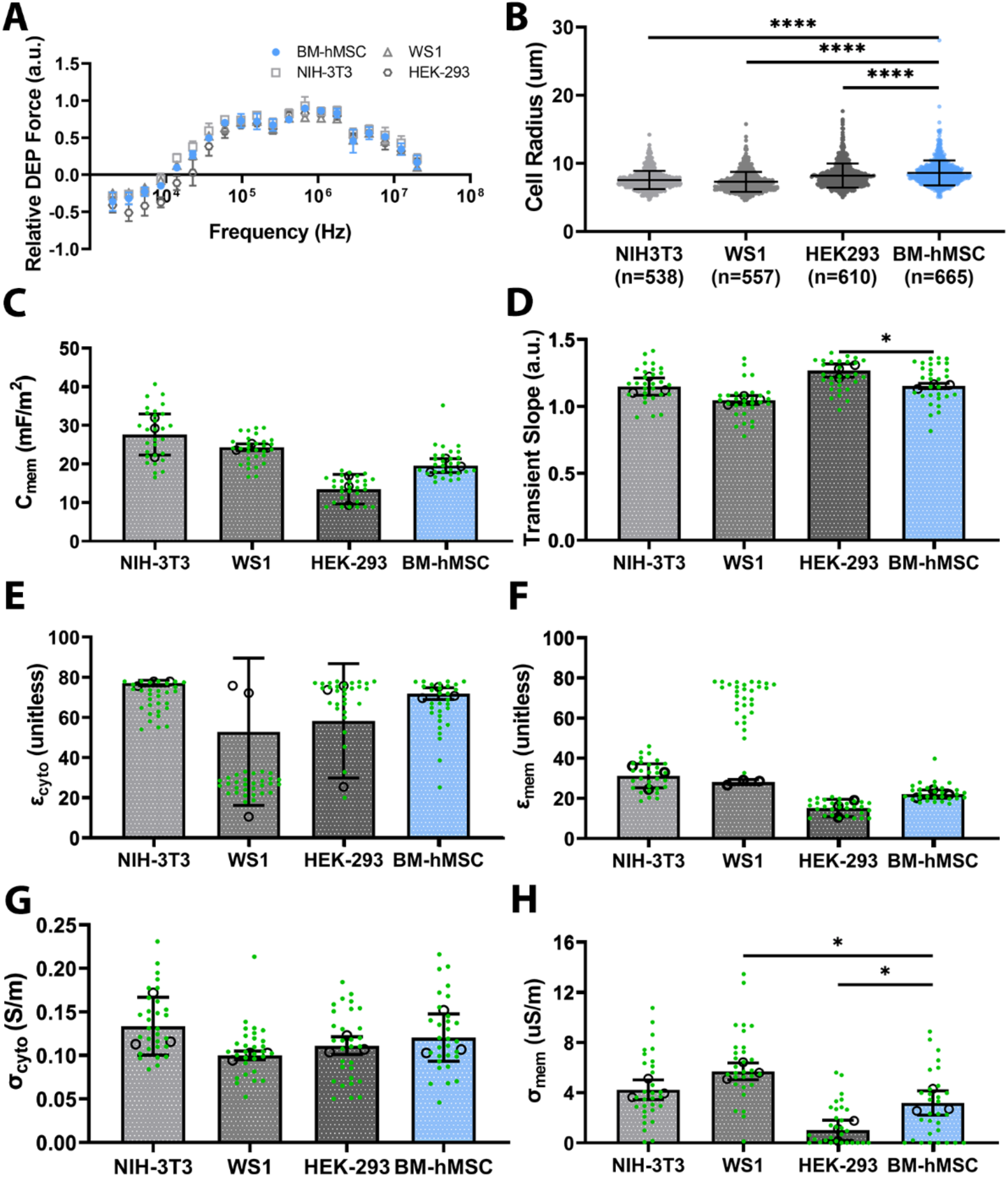
DEP cell analysis. (A) Average DEP spectrum, (B) cell size, (C) *C_mem_*, (D) transient slope, (E) *ε_cyto_*, (F) *ε_mem_* (G) *σ_cyto_*, (H) *σ_mem_* of BM-hMSCs, NIH-3T3, WS1, and HEK-293. Non-linear fitting to each individual DEP measurements (solid green circles) and non-linear fitting to pooled DEP measurements (open black circles and bars) used to estimate the electrical properties. Statistical analysis completed on pooled data sets. n = 3; *<0.05, ****<0.0001.

Comparing the two fitting methods, individual run fitting produces a larger spread than pooled fitting. The *C_mem_* of BM-hMSCs is smaller than NIH-3T3 (p=0.06) and WS1 (p=0.33) and larger than HEK-293 (p=0.17), Figure 3C. The p-values from statistical analysis is reported in parentheses; the BM-hMSCs and HEK-293 are close to significance. In Figure 3D, the transient slope of BM-hMSCs is lower than HEK-293 (*<0.05) but higher than NIH-3T3 and WS1. The *ε_cyto_, ε_mem_*, and *σ_cyto_* of BM-hMSCs is not discernible from the other cells, Figure 3E-G. There is a large variance in *ε_cyto_* for WS1 and HEK-293, possibly due to the frequency range tested, >20 MHz may provide additional information to more accurately estimate *ε_cyto_* for these cells. The *σ_cyto_* of BM-hMSCs is statistically significant compared to HEK-293 and WS1 and has a similar trend to *C_mem_*. All statistical analysis was completed on pooled data sets. Moving forward, pooled non-linear fitting will be used to estimate cell electrical properties.

### DEP and biological heterogeneity assessment of BM-hMSCs and AD-hMSCs

Transplantation studies use BM-hMSCs and AD-hMSCs, so we compared the DEP profiles of these cells, Figure 4. Physically, AD-hMSCs are larger than BM-hMSCs with an average radius 8.97±1.89 μm and 8.59 ±1.82 μm (p = 0.0003), respectively (Supplemental Figure S1). The average DEP force spectra of BM-hMSCs and AD-hMSCs is given in Figure 4A. At lower frequencies (<10^4^ Hz) the AD-hMSCs experience weaker negative DEP force compared to BM-hMSCs. The cells transition from negative to positive DEP at similar frequencies, 10^4^-10^5^ Hz. At higher frequencies (>10^5^ Hz), BM-hMSCs experienced stronger positive DEP forces than AD-hMSCs. The electrical properties were estimated and BM-hMSC and AD-hMSC have similar *C_mem_* (Figure 4B), *ε_cyto_* (Figure 4E), and *ε_mem_* (Figure 4G). The transient slope, rate of transition from negative to positive DEP, of BM-hMSC is higher than AD-hMSC with statistical significance. BM-hMSCs have a higher *σ_cyto_*, Figure 4D, while AD-hMSCs have higher *σ_mem_*, Figure 4F.

**Figure 4.**
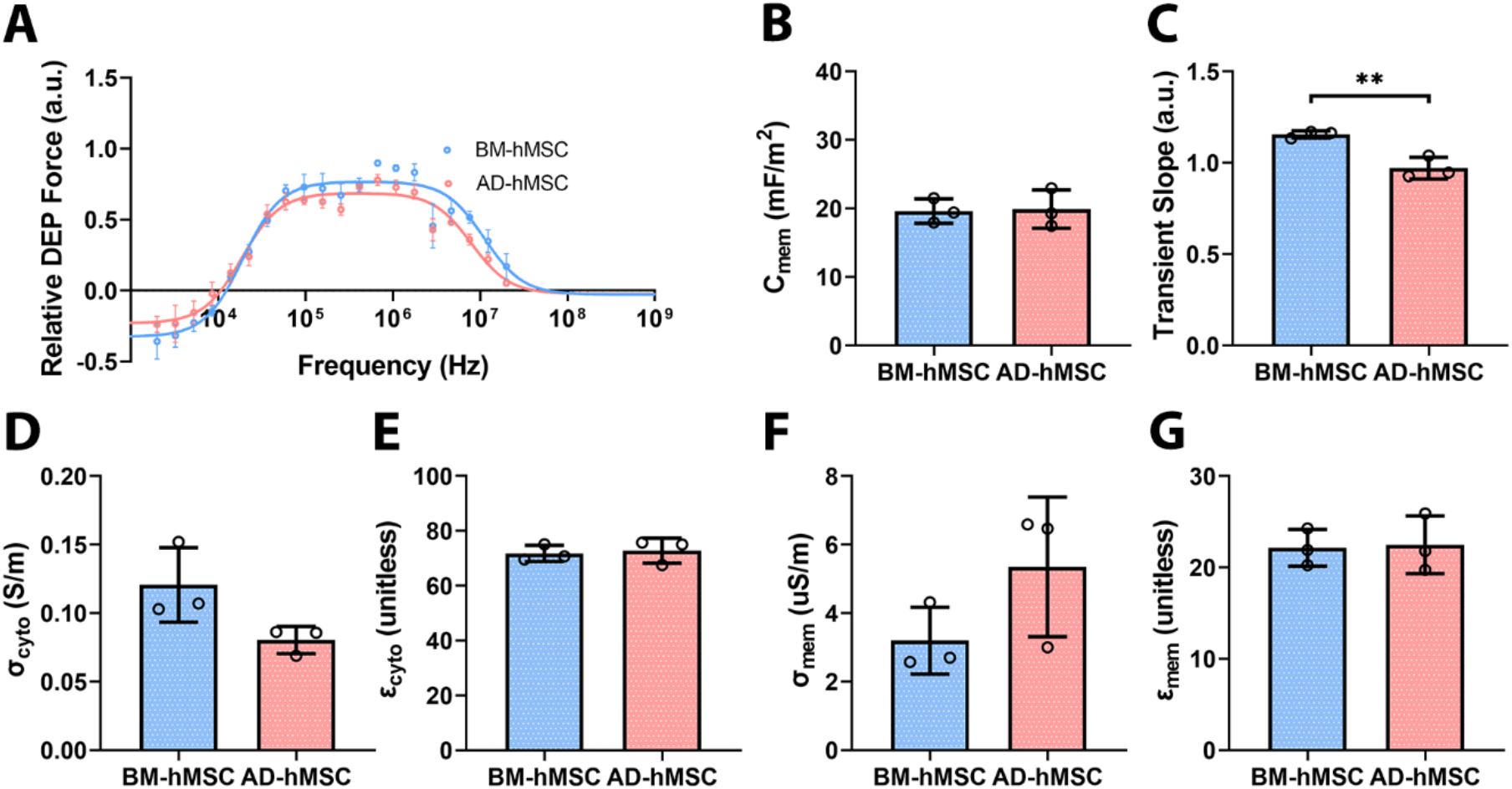
DEP Analysis of BM-hMSC and AD-hMSCs using (A) Average DEP spectrum, (B) *C_mem_*, (C) transient slope, (D) *σ_cyto_*, (E) *ε_cyto_*, (F) *σmem*, and (G) *εmem*o Non-linear fitting to pooled DEP measurements used to estimate electrical properties. n = 3; **<0.01.

Lastly, to relate the electrical properties of BM-hMSCs and AD-hMSCs back to their biological differences, we differentiated them into adipogenic and osteogenic lineages. Both BM-hMSCs and AD-hMSCs differentiated into adipocyte and osteocyte as shown by ARS staining of hydroxyapatite deposit and ORO staining of fat deposits (Figure 5A). Subtle differences in the histological data were observed. The osteocyte differentiated BM-hMSCs appear to align themselves in vertical fiber-like structure in Figure 5A while the AD-hMSCs do not. Also, the osteocyte differentiation of BM-hMSCs appear to have less calcium deposits than AD-hMSCs. Adipocyte differentiation was visualized using ORO staining in Figure 5A (bottom images) and showed that AD-hMSC has slightly more adipocyte differentiation than BM-hMSC. Gene expression data is given in Figure 5B-D, markers of adipocyte (FABP4, PPPARG, ADIPOQ) and osteocyte differentiation (ALPL, RUNX2, COL1A1) were examined. The average upregulation of ALPL, FABP4, and PPARG is higher for BM-hMSCs while both BM-hMSCs and AD-hMSCs have similar average upregulation of RUNX2. The timeline of COL1A1 shows that BM-hMSCs initially upregulate their expression (day 2-4) and then downregulate (day 6-16). Contrastingly, AD-hMSCs upregulate their expression day 6-16. The timeline of ADIPOQ shows that AD-hMSCs have negligible upregulation after day 4 whereas BM-hMSCs have a 100-fold increase from day 4 to day 16.

**Figure 5.**
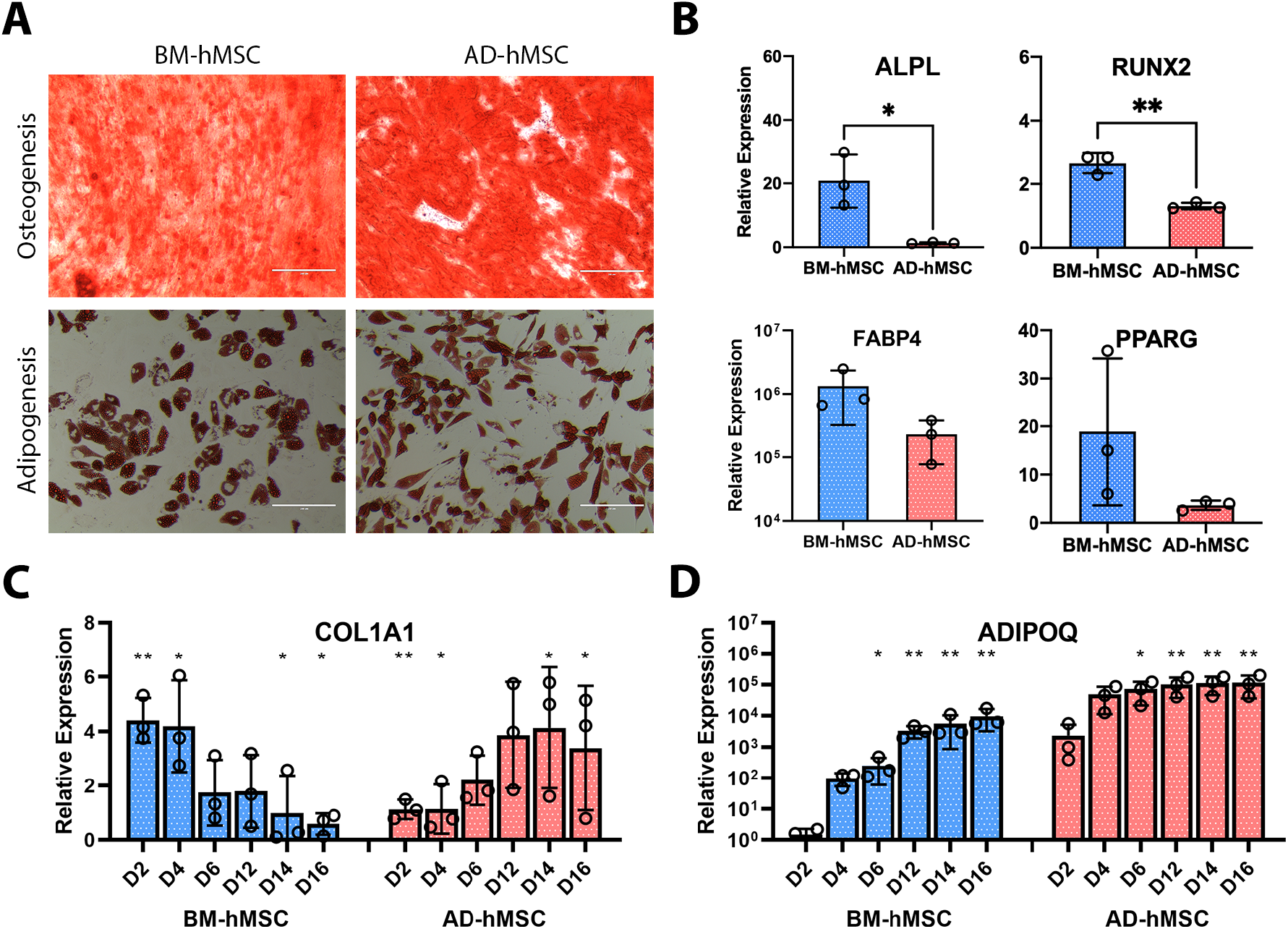
Differentiation of BM-hMSC and AD-hMSCs. (A) Osteogenesis assessed with Alizarin Red S (top) and adipogenesis assessed with Oil Red O (bottom). Gene expression profiles (B,C) for osteogenesis (ALPL, RUNX2, COL1A1) and adipogenesis (FABP4, PPARG, ADIPOQ). n = 3; *<0.05, **<0.01. All statistical test between BM-hMSCs and AD-hMSCs at the same day.

### Interpretation of heterogeneity assessments

Assessing the heterogeneity of hMSCs is important for transplantation therapy and DEP is a useful analytical tool that aids the quantification of cell electrical properties. Visualization of hMSCs DEP behavior at single frequencies serves as a good initial heterogeneity screening. Our results indicate the presence of at least two subtypes of BM-hMSCs (Figure 1). As a secondary heterogeneity screening, we analyzed the DEP profile of BM-hMSCs against NIH-3T3, WS1, and HEK-293 (suspected homogeneous cell populations). BM-hMSCs’ DEP profile indicates they are similar to NIH-3T3 and WS1 cells and different from HEK-293 cells (Figure 3). This difference may be explained using DEP theory. When cells are subjected to low frequencies the electric field interrogates the membrane properties ^[27]^, BM-hMSCs, NIH-3T3, and WS1 have fibroblastic morphology suggestive of similar protein folding and HEK-293 cells have epithelial morphology suggestive of most dissimilar protein folding. Transient slope indicates that BM-hMSCs are more heterogeneous than HEK-293 cells while WS1 are more heterogeneous than BM-hMSCs. There is literature evidence supporting that WS1 cells are heterogeneous ^[31]^. Thus, our DEP measurements may be detecting heterogeneity based on differences in protein folding, which is supported in the literature ^[32]^.

Our modeling method pooled 10 separate DEP measurements yielding non-linear curve fitting to 200 data points versus 20 data points. From the secondary assessment of hMSCs heterogeneity *σ_mem_* has a strong inverse correlation to transient slope (R-value = 0.96). BM-hMSCs and AD-hMSCs were assessed side-by-side and their DEP profiles highlights differences in heterogeneity. They have different cytoplasm and transient slope properties (Figure 4). We know from previous data that transient slope reflects heterogeneity^[33]^. Accordingly, the transient slopes here indicates that BM-hMSCs and AD-hMSCs have different subsets of cells present, while *σ_cyto_* distinguishes the sources of hMSCs.

The gene expression profiles showed that transcriptomic changes upon differentiation are not identical between BM-hMSCs and AD-hMSCs. Both sets of cells differentiated into osteocytes and adipocytes (Figure 5). ALPL^[34, 35]^, RUNX2^[36]^ and COL1A1^[37]^ are early markers of osteogenesis, and they are expected to upregulate then down regulate during osteocyte differentiation. Additionally, RUNX2 is a “master” transcriptional regulator of osteogenesis^[36]^ meaning it controls ALPL and COL1A1 expression. Overall, it takes AD-hMSCs longer to upregulate ALPL, RUNX2 and COL1A1 (Figure 5B-C and Supplemental Figure S2). Thus, the BM-hMSCs have higher osteogenic differentiation potential. PPARG, master regulator of adipogenesis^[38]^, is an early marker of adipogenesis^[39]^ and is upregulated more in BM-hMSCs (Figure 5B and Supplemental Figure S3). FABP4^[40]^ and ADIPOQ ^[41, 42]^ are late markers of adipogenesis and have similar expression patterns in the hMSCs (Supplemental Figure S3 and Figure 5D). FABP4 is upregulated more in BM-hMSCs whereas ADIPOQ is upregulated more in AD-hMSCs. Thus, the BM-hMSCs have higher adipogenic differentiation potential.

Our DEP and differentiation profiles illustrate the differences in heterogeneity between different sources of hMSCs and the BM-hMSCs have higher differential potential in comparison to AD-hMSCs. Thus, the DEP spectra, transient slope, *σ_mem_*, and *σ_cyto_* are good metrics to discern cell types and assess heterogeneity of different sources of hMSCs.

## CONCLUSIONS

Populations of hMSCs from different sources vary in heterogeneity. We show here that DEP is a good tool to assess cellular heterogeneity. The DEP profile correlates well with the differentiation profile of the BM-hMSCs and AD-hMSCs emphasizing differences in their differentiation potential. It is important to use the multiple metrics from the DEP profile (spectra, transient slope, *σ_mem_* and *σ_cyto_*) in the assessment of heterogeneity. This information will be synthesized to design DEP-based cell sorting strategies for BM-hMSCs and AD-hMSCs to selectively enrich osteocyte-biased and adipocyte-biased cell populations. Producing purified subsets of human stem cells in a label-free manner is important for screening their biological function and the advancement of stem cell therapies.

## ACKNOWLEDGEMENTS

This work was supported by the National Science Foundation CAREER award via CEBT 2048221.

## CONFLICT OF INTEREST

The authors have declared no conflict of interest.

## Notes

### Competing Interest Statement

The authors have declared no competing interest.

