## Supplemental Information for "Electrical signature of heterogeneous human mesenchymal stem cells"

The average cell size of all the of adipose-derived (AD) hMSCs was measured and plotted with the size of the NIH-3T3, WS1, HEK-293, and BM-hMSCs. Each population of cells differ in size with NIH-3T3 < WS1 < HEK-293 < BM-hMSCs < AD-hMSCs.

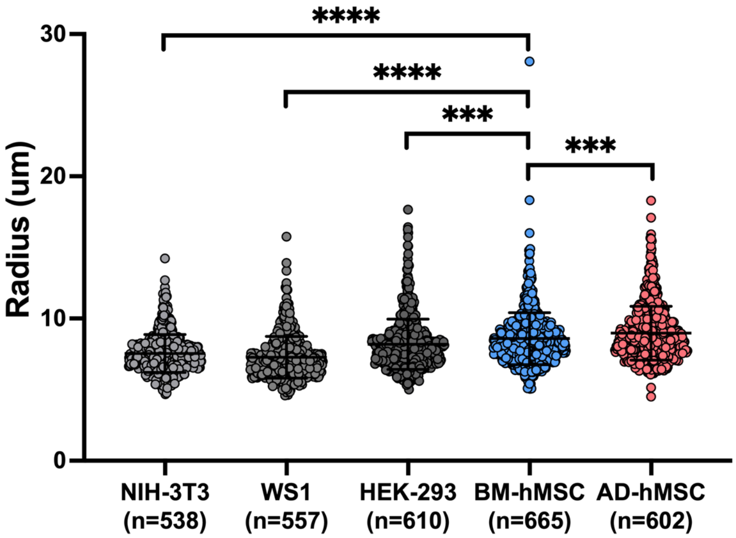


**Figure S1.** Radius comparison for all cell types. The distribution of cell radiuses quantified using ImageJ. The images were obtained using trypan blue exclusion under a 20x phase contrast objective. The "n" reported under the type of cells is the number of cells measured. n = 3 individual experiments; ***<0.001, ****<0.0001.

The relative gene expression of ALPL and RUNX2 (osteogenic markers) over time is given in Figure S2.A-B. Overall for both the BM-hMSCs and AD-hMSCs ALPL upregulates from day 0 to day 16 with the magnitude of upregulation being higher for BM-hMSCs. RUNX2 initially upregulates then down regulates around day 4 and day 10 for the BM-hMSCs and the AD-hMSCs, respectively. The magnitude of upregulation is higher for the BM-hMSCs. From both gene expression assessments, it can be seen that BM-hMSCs have an easier time turning into osteocytes.

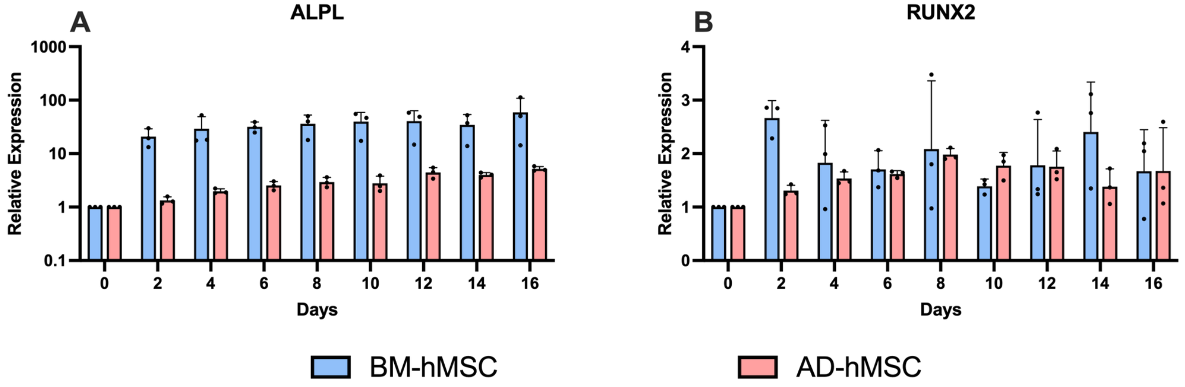


**Figure S2.** Time-course gene expression profiles of (A) ALPL and (B) RUNX2 for the osteogenic differentiation of BM-hMSCs and AD-hMSCs. Error bars shown are standard deviation calculated from repeat experiments (n=3).

The relative gene expression of ADIPOQ, FABP4, and PPARG (adipogenic markers) over time is given in Figure S3.A-C. Overall these markers upregulate for the BM-hMSCs and AD-hMSCs from day 0 to day 16. ADIPOQ is upregulated more in AD-hMSCs while PPARG is upregulated more in BM-hMSCs. In the case of FABP4 it is slightly upregulated more in the BM-hMSCs. From these results, the BM-hMSCs have a higher adipogenic potential.

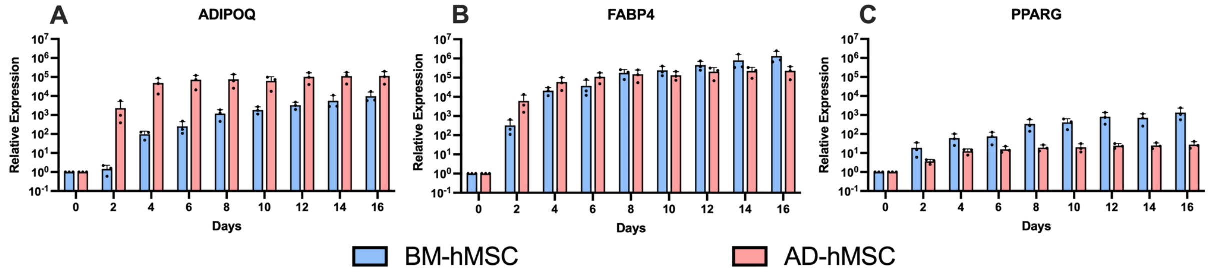


**Figure S3.** Time-course gene expression profiles of (A) ADIPOQ, (B) FABP4, and (C) PPARG for the adipogenic differentiation of BM-hMSCs and AD-hMSCs. Error bars shown are standard deviation calculated from repeat experiments (n=3).

The primer sequences are provided in the Table S1.

**Table S1.** Primer Information

| Target | FWD Primer 5' - 3' | REV Primer 5' - 3' | IDT Assay ID |
|---|---|---|---|
| GAPDH | GACAGTCAGCCGCATCTTCT | GCGCCCAATACGACCAAATC | - |
| ALPL | - | - | Hs.PT.56a.40555206 |
| RUNX2 | - | - | Hs.PT.56a.19568141 |
| COL1A1 | CCCTCCACTCCTTCCCAAATC | CTTCCTGACTCTCCTCCGAAC | - |
| FABP4 | CATAACCTTAGATGGGGGTGTCC | GTCCCTTGGCTTATGCTCTCTC | - |
| PPARG | GCTTGTGAAGGATGCAAGGG | ATCCGCCCAAACCTGATGG | - |
| ADIPOQ | TTCCGCAGTGTAGGCTTTACC | GTGTGGCTTGGGGATACGAG | - |

The Forward and reverse primer pair information for qPCR gene expression assay. In the case where a commercially available primer pairs were used; the assay ID is supplied in place of the actual primer sequences.
